# Towards a bottom-up understanding of antimicrobial use and resistance on the farm: A knowledge, attitudes, and practices survey across livestock systems in five African countries

**DOI:** 10.1101/703298

**Authors:** Mark A Caudell, Alejandro Dorado-Garcia, Suzanne Eckford, Denis Byarugaba, Kofi Afakye, Tamara Chansa-Kabali, Emmanuel Kabali, Stella Kiambi, Tabitha Kimani, Geoffrey Mainda, Peter Mangesho, Francis Chimpangu, Kululeko Dube, Bashiru Boi Kikimoto, Eric Koka, Tendai Mugara, Samuel Swiswa

## Abstract

The nutritional and economic potentials of livestock systems are compromised by the emergence and spread of antimicrobial resistance. A major driver of resistance is the misuse and abuse of antimicrobial drugs. The likelihood of misuse may be elevated in low- and middle-income countries where limited professional veterinary services and laissez faire access to drugs are assumed to promote non-prudent practices (e.g., self-administration of drugs). The extent of these practices, as well as the knowledge and attitudes motivating them, are largely unknown within most agricultural communities in low- and middle-income countries. The main objective of this study was to document dimensions of knowledge, attitudes and practices related to antimicrobial use and antimicrobial resistance in livestock systems and identify the livelihood factors associated with these dimensions. A mixed-methods ethnographic approach was used to survey households keeping layers in Ghana (N=110) and Kenya (N=76), pastoralists keeping cattle, sheep, and goats in Tanzania (N=195), and broiler farmers in Zambia (N=198), and Zimbabwe (N=298). Across countries, we find that it is individuals who live or work at the farm who draw upon their knowledge and experiences to make decisions regarding antimicrobial use and related practices. Input from animal health professionals is rare and antimicrobials are sourced at local, privately owned agrovet drug shops. We also find that knowledge, attitudes, and particularly practices significantly varied across countries, with poultry farmers holding more knowledge, desirable attitudes, and prudent practices compared to pastoralists households. Multivariate models showed that variation is related to several factors, including education, disease dynamics on the farm, and sources of animal health information. Study results emphasize that interventions to limit antimicrobial resistance must be founded upon a bottom-up understanding of antimicrobial use at the farm-level given limited input from animal health professionals and under-resourced regulatory capacities within most low- and middle-income countries. Establishing this bottom-up understanding across cultures and production systems will inform the development and implementation of the behavioral change interventions to combat AMR globally.

## Introduction

Antimicrobial drugs are essential to maintain animal health within livestock production systems but their misuse and/or abuse increases selection for the emergence, transmission, and persistence of antimicrobial resistance (AMR) [1–3]. AMR results in therapeutic failures and increases lengths and/or cycles of treatment, thereby producing downstream impacts on animal welfare, food security, and public health. AMR in animals may impact prevalence in people as resistant microorganisms (e.g., bacteria, viruses, parasites) can be transmitted across the human-animal interface through a diverse set of pathways, including consumption of animal products, direct contact and sharing of water sources [2,4]. This potential for transmission is significant considering nine of the fourteen classes of drugs labelled ‘critically important’ in public health are also used in livestock systems [5]. Antimicrobial use in the agricultural sector is projected to increase by 67% by the year 2030, potentially further compromising their effectiveness in both animal and human health [6]. The interconnectedness of antimicrobial use in agriculture and public health has led to AMR being referred to as the “quintessential One Health issue” of our time [7].

One Health implications of AMR are particularly relevant in low- and middle-income countries (LMICs) where high burdens of infectious disease often exist alongside livelihoods that promote frequent interactions between people and livestock. Genotypic studies in LMICs and high-income countries (HICs) show these contexts can promote AMR transmission between people, animals and the environment. In Tanzania, for example, genotypic similarity has been documented between resistant enteric bacteria in people, animals (both livestock and wildlife) and the environment (i.e., waters sources) [8,9] Similar patterns were found in *Salmonella* isolates from people and animals in Uganda [10]. In contrast, genotypic studies from high-income countries have largely shown distinguishable epidemics of AMR in livestock and the general population [11–13]. These patterns within HICs are consistent with limited contact between the general population and livestock and with the more developed health, sanitation, and regulatory infrastructures that limit transmission events [11–15]. While a study conducted within Netherlands did document overlap between livestock (pigs) and people, the overlap was dependent on intensity of contact and increased in farming communities [16]. These studies demonstrate that intervention efforts to limit AMR will diverge globally and must be adapted to national and local realities.

Within LMICs, the combined realities of underfunded veterinary healthcare systems and limited regulatory capacities often constrain efforts to promote prudent AMU and limit AMR in the agricultural sector [17–20]. Increasingly, these efforts will be guided by multisectoral National Action Plans (NAPs) whose activities are supported by the Tripartite Collaboration on AMR consisting of the Food and Agriculture Organization (FAO), the World Organization for Animal Health (OIE), and the World Health Organization (WHO). NAPs set a series of goals to improve awareness on AMR and related threats, develop capacity for surveillance and monitoring of AMU/AMR, strengthen governance, and promote AMU stewardship within the public and animal health sectors [21,22]. In many LMICs, implementation of these NAPs within the livestock sector is constrained by limited access to animal health professionals (see Figure 1). In the five African countries surveyed in this study, for example, the ratio of veterinarians to livestock is about 20 times lower compared to high-income countries of Denmark, France, Spain and the USA (data obtained from WAHIS and FAO-STAT). Accessibility issues are common across LMICs given private sector healthcare services have not adequately compensated for the reductions in public services forced by Structural Adjustment Programs in the 1980’s [23,24]. Further limiting achievement of NAP goals are under resourced regulatory authorities (e.g., national medical authorities, food safety departments), which make national regulations, such as laws mandating prescriptions for antimicrobials or regulations on antibiotic residues, difficult to enforce [20].

**Figure 1.**
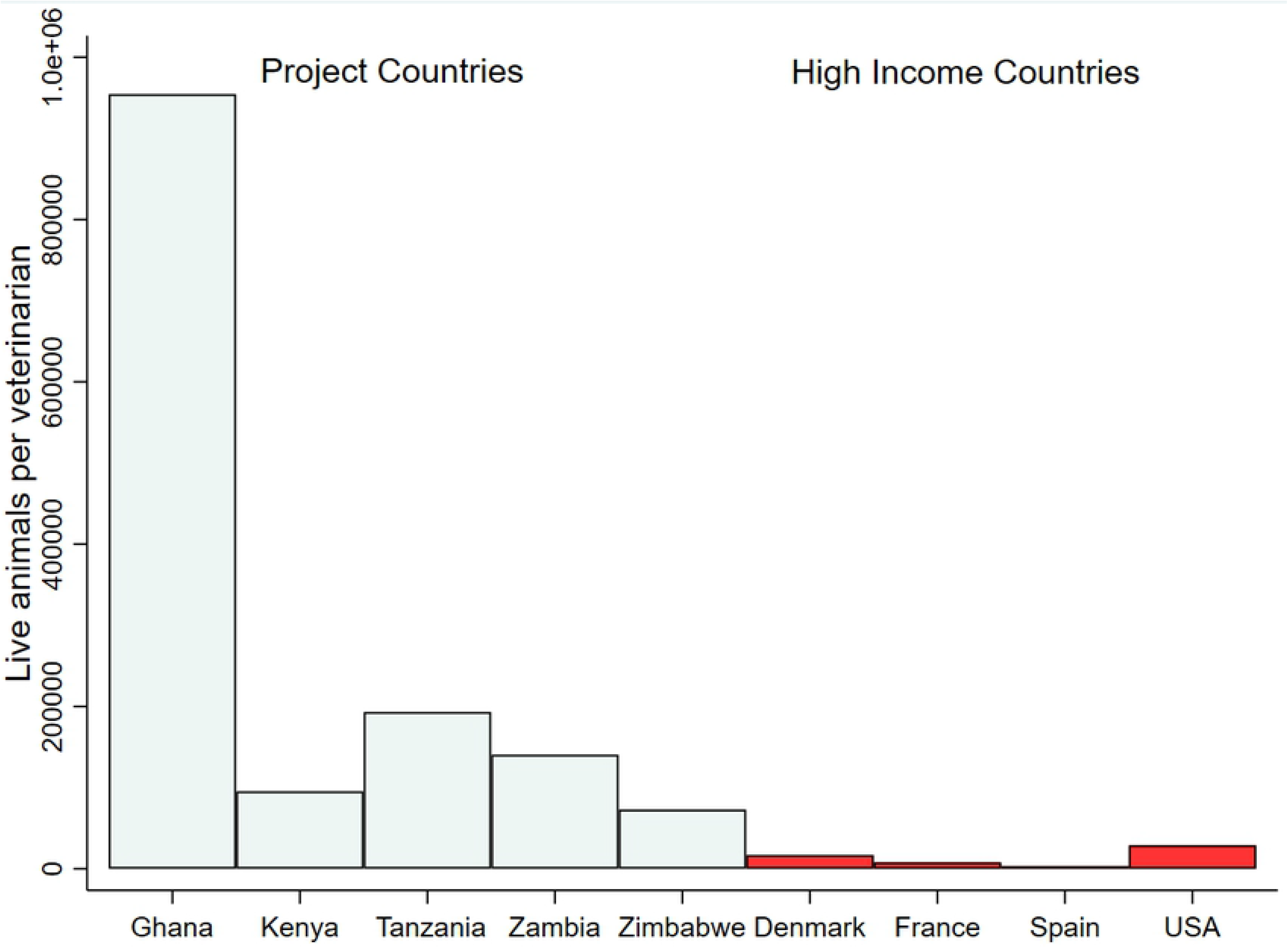
Live animals per veterinarian for project countries compared with High Income Countries. Data obtained from WAHIS and FAO-STAT.

Constraints on the resources that are needed to support top-down implementation and enforcement of NAP strategies within LMICs emphasize the need for behavioral change interventions targeted at the household level. These “bottom-up” intervention approaches will be informed by understanding the sociocultural, economic, and historical factors that motivate antimicrobial use and related practices (e.g., observance of withdrawal) in livestock systems. Unfortunately, within most LMICs, there is currently little information on AMU practices and motivating factors in livestock systems [2,19,25]. Available studies generally find that farmers administer antimicrobials themselves, mostly without prescriptions or using input from animal health professionals, and often engage in other non-prudent practices, such as not observing withdrawal periods [26–35]. However, there has not been a comprehensive study of the knowledge, attitudes, and practices (KAP) associated with AMU or AMR to identify the factors associated with observance of prudent or non-prudent AMU practices. Lacking knowledge of KAP and associated factors hinders the development and implementation of targeted behavioral change interventions to promote prudent AMU and limit AMR across livestock systems in LMICs. To inform the development of these interventions, we conducted a KAP survey in livestock systems in five African countries. The systems surveyed included pastoralist communities in Tanzania, large-scale and intensive commercial poultry farmers in Ghana (layers) and small-scale commercial poultry farmers in Kenya (layers), Zambia (broilers), and Zimbabwe (broilers).

## Materials and Methods

### Study Locations

The KAP survey was administered to 887 farmers in Ghana, Kenya, Tanzania, Zambia, and Zimbabwe. Three types of production systems were targeted, including broilers (Zambia and Zimbabwe), layers (Ghana, and Kenya,) and pastoralists (Tanzania). Production systems were chosen considering several factors, including contribution to national agricultural GDP, the projected growth of the system, the patterns of AMU generally associated with the system, and input from governmental partners (e.g., ministries of livestock). Geographic areas targeted for survey were chosen based upon census records of the number of households pursuing the selected system, if available, and/or through discussions with local animal health officers. Figure 2 provides a map of the countries surveyed, the approximate survey locations within each country, the production system surveyed, and corresponding sample size for each country.

**Figure 2.**
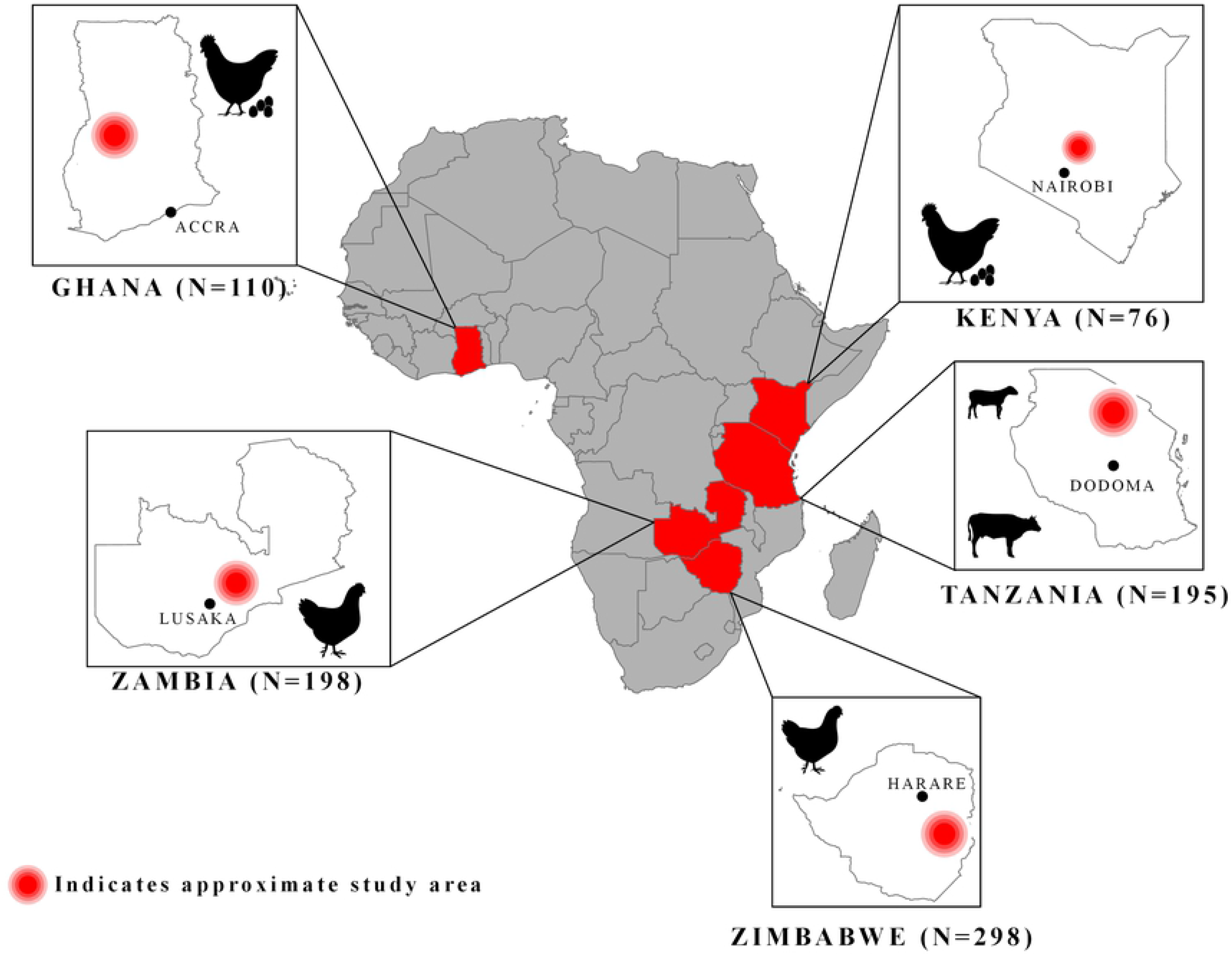
Project map. Surveyed countries are highlighted in red. Insets of country maps include country capitals, approximate study area (red circle), icons for production system surveyed, and sample size below.

### KAP Survey Development and Deployment

The project was developed and implemented by an interdisciplinary research team comprised of animal health experts, epidemiologists and social scientists from FAO, national ministries of agriculture and livestock, and within-country academic institutions. The study employed a modified version of the exploratory cross-sectional survey design [36] using qualitative data to inform quantitative surveys and confirm quantitative measures. Focus Group Discussions (FGDs) and Key Informant Interviews (KIIs) were conducted with stakeholders influencing AMU/AMR patterns, including farm owners, farm managers, and farm workers, animal health professionals (e.g., district veterinary officers, community animal health workers, para-veterinarians), owners and employees of shops selling antimicrobials and other livestock products (e.g., feed, disinfectant, vitamins) and feed producers and distributors in poultry systems. When possible, we attempted to interview a minimum of three to a maximum of six groups within each stakeholder category. Three to six groups were targeted given that 90% of the information within a research topic is normally discoverable with this range (i.e., the saturation point of a topic) [37].

Qualitative interviews with stakeholders were concentrated around major themes associated with AMU and AMR including farm management practices, local disease histories, health-seeking behaviors and health infrastructures, and governance, regulations, policies, and enforcement related to AMU and AMR. Thematic analysis of qualitative data was used to develop a KAP survey instrument of over 200 items (see S2: KAP questionnaire). KAP questionnaires were administered by local enumerators using the Kobo Collect^®^ application. Enumerators received three to four days of training and assisted in refining the survey as most had an animal health background and worked in areas near the study communities. Surveys were piloted prior to administration to ensure question clarity, conduct further refinement, and verify survey times. Across the five project countries, surveys were completed between 45 minutes and 1 hour and 30 minutes. All informants provided consent to participate through signature or thumbprint.

Several sampling strategies were used across project countries given differences in availability of census records, reachability of households, financial resources, and time constraints. In most countries, a purposive sampling approach was used given a lack of information on the households that participated in a particular production system (e.g., the number of households in a community that kept broilers). In Kenya and Zambia, enumerators developed lists of households keeping layers or broilers, respectively, by walking house-to-house and/or consulting with local leaders and veterinary officers. These sample frames were then consulted to schedule interviews. In Ghana and Zimbabwe, existing lists of farmers were used alongside a snowball sampling approach where farmers identified other poultry farmers within the community. For Tanzania, two selection strategies were used including visiting the Maasai households and at market places. See S1: Study Locations and sampling for more country-specific information on sampling criteria and areas and populations surveyed.

### Ethical Approvals

Ethical approvals were received in each country. For Ghana, the study was approved by the Ministry of Health Ethics Review Committee (ID No. 014/10/18). For Kenya, the study was approved by the AMREF Health Africa Ethics and Scientific Review Committee (AMREF-ESRC P551/2018) and the Institutional Animal Care and Use Committee of KALRO-Veterinary Science Research Institute (KALRO-VSRI/IACUC016/28092018). For Tanzania, the study was reviewed and approved by the Medical Research Coordinating Committee of the National Institute for Medical Research (NIMR) in Tanzania and certificate clearance no. NIMR/HQ/R.8a/Vol.IX/2926 was issued. For Zambia, the study was reviewed and approved by the ERES Converge Ethic Committee, Ref: No 2018-Nov-020 was issued. For Zimbabwe, approval was obtained from the Agriculture Research Council (ARC) of the Department of Veterinary Services & Crop and Livestock (Reference Number: 008/2018).

### Variable Description and Data Analysis

Linear scales of knowledge, attitudes and practices were generated from the variables included in Table 1 and used as outcomes in ordinal least squares (OLS) regression models. Scales were developed by recoding variable answers to binary with 1 representing sufficient knowledge, desirable attitudes and appropriate practices for control of AMR and 0 representing insufficient knowledge, undesirable attitudes and inappropriate practices. For attitudes and practices scales, responses of “indifferent” and “sometimes” were coded undesirable and inappropriate, respectively. Responses were then summed for each participant and divided by the total number of items within the category to arrive at a percentage of correct answers. For example, if a respondent reported observing 6 of the 8 appropriate practices they received a 75% (6/8) prudent practice score out of a possible 100%. One-Way ANOVA analysis were used to assess significant differences in KAP scores across countries. Tukey-Kramer comparisons were used to assess which country comparisons were significant. Pairwise correlations were used to calculate the associations between KAP scores across and within countries.

**Table 1.** Description of variables included in knowledge, attitudes, and practices scale.

| Original variables | Recoded values |
| --- | --- |
| **Knowledge (Yes=1 No=0)** |  |
| Able to explain antimicrobial resistance | Yes =1; No=0 |
| Able to explain what antibiotics are/do | Yes =1; No=0 |
| Able to explain what antibiotic residues are/do | Yes =1; No=0 |
| Able to explain dosage/treatment of commonly used antibiotic | Yes =1; No=0 |
| Able to explain what vaccines are/do | Yes =1; No=0 |
| **Attitudes (disagree =1, neutral/indifferent = 2, agree=3)** |  |
| If medicines are given TOO OFTEN then they might stop working | 1&2=0; 3=1 |
| Giving animals that are not sick antimicrobials will PREVENT THEM FROM BECOMING SICK in the future | 1=1; 2&3=0 |
| Giving animals antimicrobials can help them GROW BIGGER, FASTER, FATTER, boost egg production/size | 1=1; 2&3=0 |
| It is important to get CONSULTATION FROM A VETERINARIAN before giving antimicrobials to the animals | 1&2=0; 3=1 |
| Using vaccines can reduce use of antibiotics | 1&2=0; 3=1 |
| After using antibiotics on an animal, you should WAIT sometime before using the PRODUCTS from it, such as bird meat/eggs/milk | 1&2=0; 3=1 |
| **Practices** (never/rarely= 0, 1= sometimes, 2=almost always) |  |
| Give antimicrobials when get you day-old chicks/new calves/smallstock | 0=1 ; 1&2=0 |
| Give antimicrobials to all animals when one is sick | 0=1 ; 1&2=0 |
| Give animals a LARGER DOSE than recommended | 0=1 ; 1&2=0 |
| Give birds a SMALLER DOSE than the recommended dose? | 0=1 ; 1&2=0 |
| STOP using antimicrobials before the full dose because the animal has improved | 0=1 ; 1&2=0 |
| Use expired medicines | 0=1; 1&2=0 |
| Have a prescription when purchasing antibiotics | 0&1=0 ; 2=1 |
| Observe withdrawal from antimicrobials | 0&1=0 ; 2=1 |

Ordinal least squares (OLS) regression was used to assess the factors associated with KAP scales. Regression models were specified that assessed three domains, including demographics, on-farm dynamics, and health seeking practices. Demographics included age, gender, education and years engaged in the targeted production system (e.g., number of years a respondent kept broilers). On-farm dynamics included variables related to farm size, biosecurity, disease histories, antimicrobial use patterns, trainings on farm management and record-keeping. The health seeking model included variables on frequency which people sought advice from various stakeholders (friends, veterinarians, agrovets, etc). Variable definitions are provided below the tables. Beyond analysis pooled across countries, separate models were also run across production systems (i.e., for poultry and pastoralist systems). In regression models that included more than one country, we control for country-level effects by entering country as dummy variables in order to assess those factors related to KAP scales across differing country contexts.

Regression diagnostics were assessed for all models. Influential data points were examined through calculating Cook’s D and rerunning models with observations associated with values exceeding 4/N (4/857=0.005). Excluding observations above this threshold did not impact model interpretation. Multicollinearity was assessed calculating variance inflation factors (VIFs). Variance inflation factors for variables of interest were less than 2, well below the recommended cut-off of 5[38]. Variance inflation factors for country-control variables, specifically Tanzania (VIF≈6) were above the cut-off. However, we do not interpret the coefficients of control variables and these controls were not collinear with variables of interest ensuring the performance of the controls was not impaired [39]. Normality of residuals was tested with the Shapiro-Wilk Test, kernel density, and standardized normal probability plots. Results from all tests indicted only slight deviations from normality. The homoscedasticity assumption (i.e., homogeneity of variance of residuals) were tested using residuals versus fitted (predicted) plots [40]. There was evidence of minor heteroscedasticity in the demographic-factors model so a Huber-White sandwich estimator was used to provide robust estimates [41].

## Results

### Descriptive Results

Here, we focus our discussion on variables used in the KAP regression models. More detailed descriptive statistics on demographics, socioeconomics, farm management practices (including antimicrobial use), and diseases reported can be found in Table S4-Table S7. Ghana farmers kept the largest flocks with a median flock size of 4054 birds (Q_1_=2000, Q_3_=9000) while Kenya farms had a median of 700 birds (Q_1_=300, Q_3_=1150). Broiler systems were smaller than layer systems with a median of 105 birds in Zambia (Q_1_=0, Q_3_=250) and 100 birds in Zimbabwe (Q_1_=25, Q_3_=150). Flock mortalities, calculated as percentage of average flock size, were highest in layer systems in Kenya (≈16%) and Ghana (≈14%) and slightly lower in broiler systems in Zambia (≈10%) and Zimbabwe (≈8%). In terms of biosecurity, a minority of farms had footbaths with Zambia farmers reporting the highest rate of footbath ownership (41%) followed by Zimbabwe (21%), Ghana (6%) and Kenya (3%). Around 70% of farms owned boots for the poultry houses except for farmers in Zambia where only 32% reported owning boots. Ninety nine percent of farmers in Ghana reported keeping farm records, followed by ≈80% of Zambian farmers, ≈70% Zimbabwean farmers, and ≈60% of Kenyan farmers. The most common records kept on the farm were sales records and flock mortalities. Medicine costs, including antimicrobials and vaccines, were the highest in Ghana (0.65 USD per bird), followed by Zimbabwe (0.31 USD), Zambia (0.23 USD) and Kenya (0.21 USD). The top three self-reported diseases in layers were Coccidiosis (≈63%), Chronic Respiratory Disease (CRD) (76%) and Newcastle Disease (39%). For broilers, the top three self-reported diseases were Coccidiosis (≈43%), CRD (≈32%), and Infectious Bronchitis (≈30%).

Although of comparable age (≈45 years), Maasai were distinguished from other project communities in terms of household size, farm owner gender, education levels, farm management training, and record-keeping. Maasai households averaged around 18 persons while poultry households averaged around 5 persons. Ninety-three percent of farm owners were men compared to ≈65% in poultry systems. Over 60% of Maasai reported having no formal education while this percentage was 0 in Zambia and Zimbabwe, 1% in Kenya and 10% in Ghana. Only 4% of Maasai households reported having training by animal health professionals or organizations (e.g. NGOs) on any aspect of farm management (record-keeping, animal health, etc.) and 10% reported keeping written records. The Maasai owned a median of 68 cattle (Q_1_=20, Q_3_=160) and 120 sheep and goats (Q_1_=60, Q_3_=255). Only 4% of Maasai households reported having training by animal health professionals or organizations (e.g. NGOs) on any aspect of farm management (record-keeping, animal health, etc.). Maasai keep these animals divided into those at the “temporary bomas” (usually distant locations to ease local grazing pressures and cope with drought), those that move in and out of the household for daily grazing, and those kept inside the household (mostly young or sick animals). A median of 30 cattle (Q_1_=9, Q_3_=60) and 0 sheep and goats (Q_1_=0, Q_3_=7) were kept in temporary bomas. A median of 30 cattle (Q_1_=11, Q_3_=70) and 79 goats (Q_1_=35, Q_3_=150) moved in and out for daily grazing. A median of 5 cattle (Q_1_=1, Q_3_=7) and 6 goats (Q_1_=1, Q_3_=8) were kept at home. The most common self-reported diseases for cattle were Contagious Bovine Pleuropneumonia (70%), Coenurosis (63%) and East Coast Fever (61%). For sheep and goats, the top three diseases were Coenurosis (96%), Contagious Caprine Pleuropneumonia (92%) and Sheep and Goat Pox (54%).

### Antimicrobial Knowledge and Use Patterns

Across countries, 104 antimicrobial products were reported as commonly used, containing active substances representing eight antimicrobial classes. 70% of these products (N=73) contain a tetracycline and 20% (N=45) contained a macrolides or aminoglycoside. The most common reason for using antibiotics was for treatment followed by preventing sickness in groups and individual animals and, mostly in the Maasai, for faster and bigger growth (Figure 3). The average number of products reported by households as commonly used was around six. Zambia and Tanzania reported the highest use of antimicrobials (≈3-5 times per month) followed by Zimbabwe and Ghana (1-2 times per month) and Kenya (<1-2 times per month). Most households acquired antibiotics from agrovet shops (≈83%) with much lower percentages (≈12% for each source) acquiring drugs from feed distributors, shops that were not agrovets (e.g., hardware stores) and government vets (see Table S8). When purchasing AMs at agrovet shops, ≈38% reported providing symptoms of sick animals and getting advice on the specific AM (≈38%), while ≈30% were told the AMs they needed but given no instructions on use and 25% told the agrovet the AMs they needed and did not receive any instructions on use (see Table S9). Very few households (12%) reported almost always having a prescription when purchasing antimicrobials. Observation of withdrawal was also limited with an average of 10% of household discarding products or meat from animals undergoing treatment or within the withdrawal period. Around 50% of farmers reported to observe an increase in treatment failures since they began farming.

**Figure 3.**
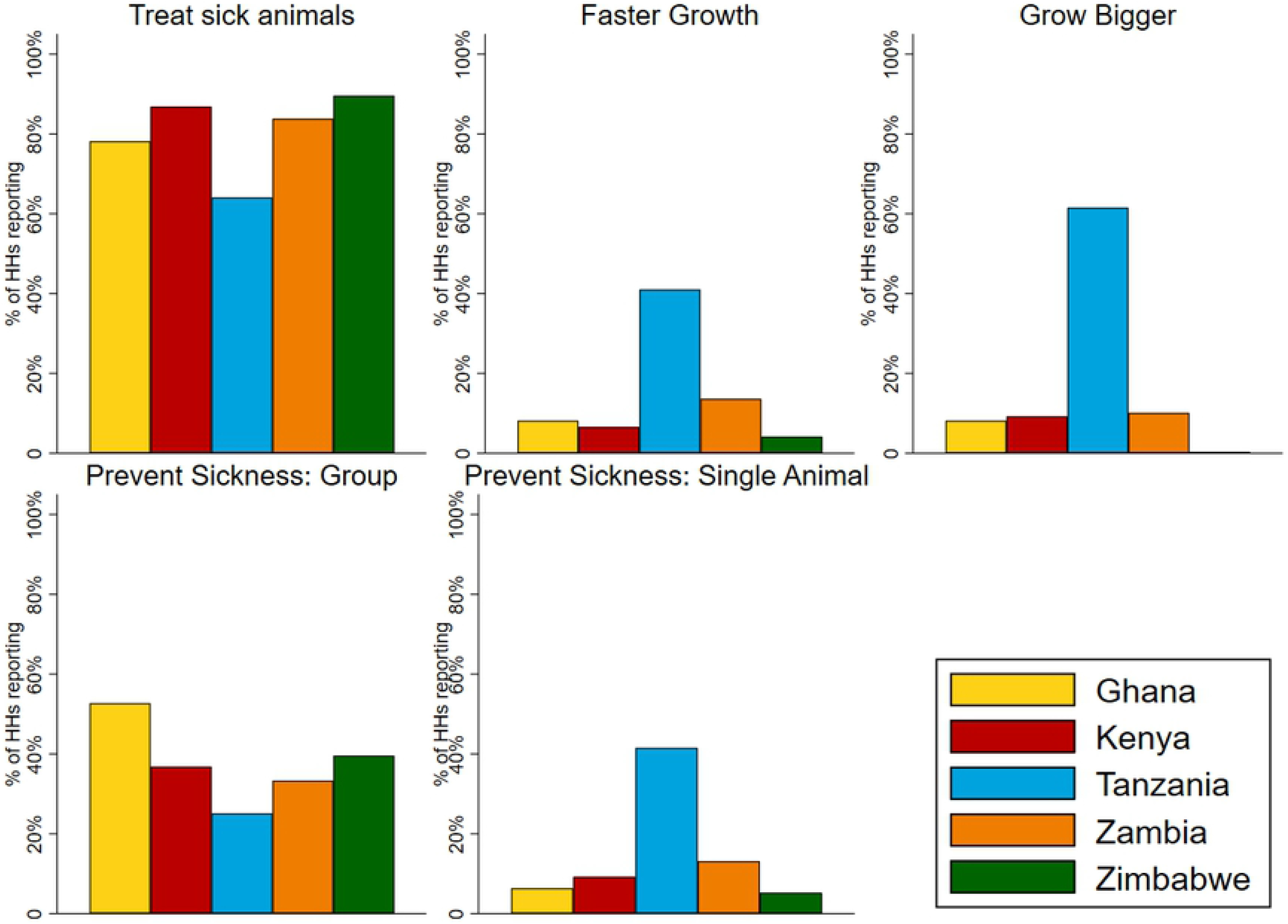
Reported reasons for using antimicrobials in livestock.

Sources of health advice related to antimicrobial use are provided in Figure 4. A majority of households reported almost never or rarely asking advice from feed distributors (75%, only asked in poultry systems), private veterinarians (74%), community health workers (63%), government veterinarians (55%) and agrovets (51%). When administering antibiotics, most respondents indicated that the farm owner almost always administered the antibiotics (53%), followed by the farm manager (24%) and family and friends (8%) (see Figure 5). Few households reported that government vets, private vets, or agrovets administered antimicrobials to their animals.

**Figure 4.**
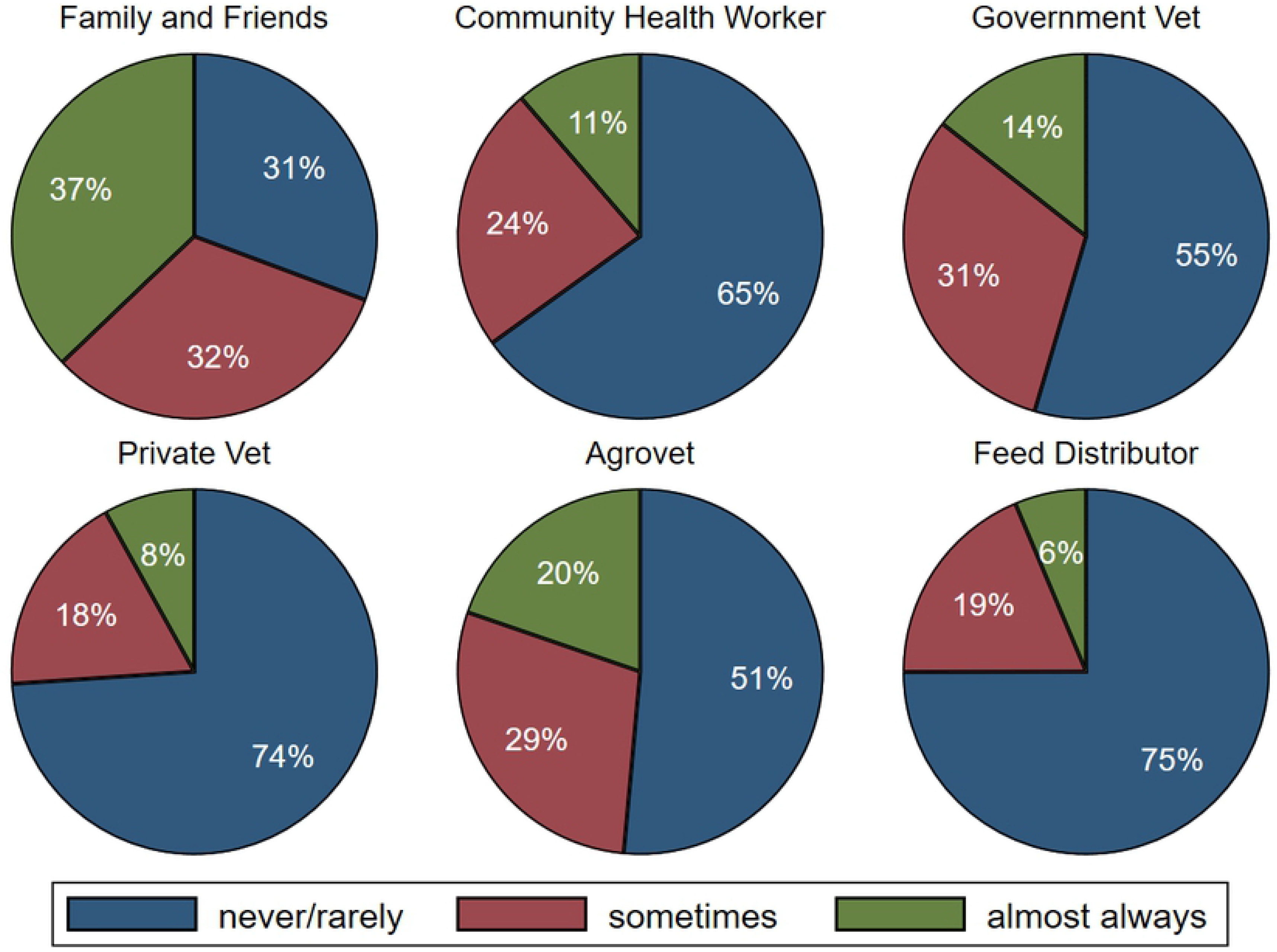
Sources of advice on animal health. Advice from feed distributor was not asked in Tanzania given the Maasai do not purchase feed for their livestock. N=867

**Figure 5.**
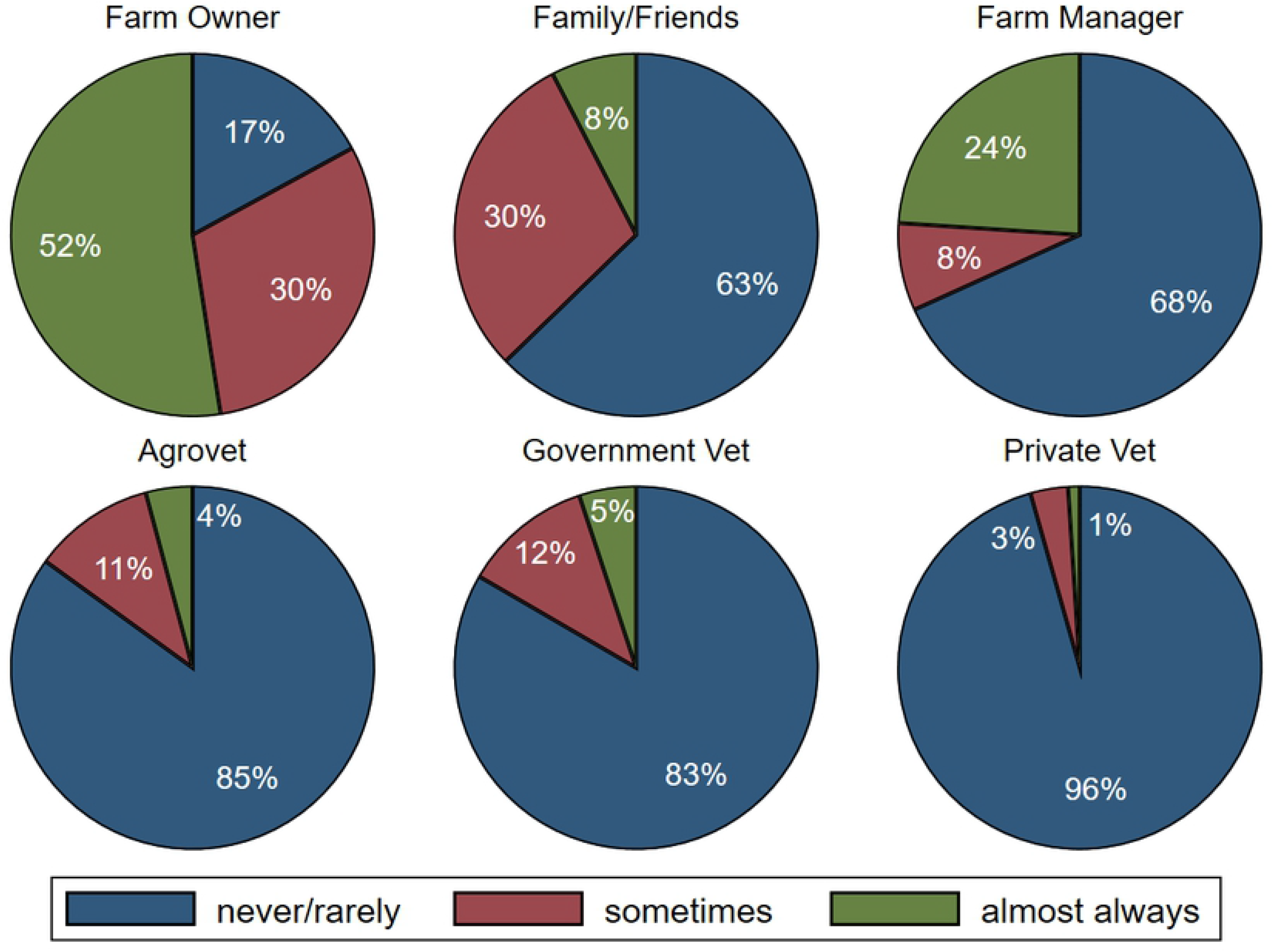
People administering antimicrobials to livestock. N=867 with the exception of Farm Manager which is based on 672 observations (Maasai do not have farm managers).

### Distribution of Knowledge, Attitudes and Practices

Figure 6 provides the distribution of the percentage of correct responses for AMU/AMR knowledge, attitudes, and practices across the project countries. One Way ANOVA showed significant differences across countries between percentages correct on AMU/AMR related knowledge (F(4,862) = 16.89, p< 0.00), attitudes (F(4,862) = 66.26, p< 0.00), and practices (F(4,862) = 159.71, p< 0.00). Tukey-Kramer comparisons demonstrated that significant differences (p<0.05) in knowledge and attitudes across countries were largely driven by differences between pastoralist households (Tanzania) and poultry production households. While the median correct AMU/AMR knowledge and attitudes were around 70% in poultry system countries, the medians for Tanzania were around 40%. In contrast to knowledge and attitudes, there were significant differences in practices across all countries, except between the broiler keeping households in Zambia and Zimbabwe, which reported the greatest adherence to prudent practices (≈85%). See for Figures S6-S8 for One Way ANOVA results and Tukey and Hamer minimum significant differences for each pair comparison.

**Figure 6.**
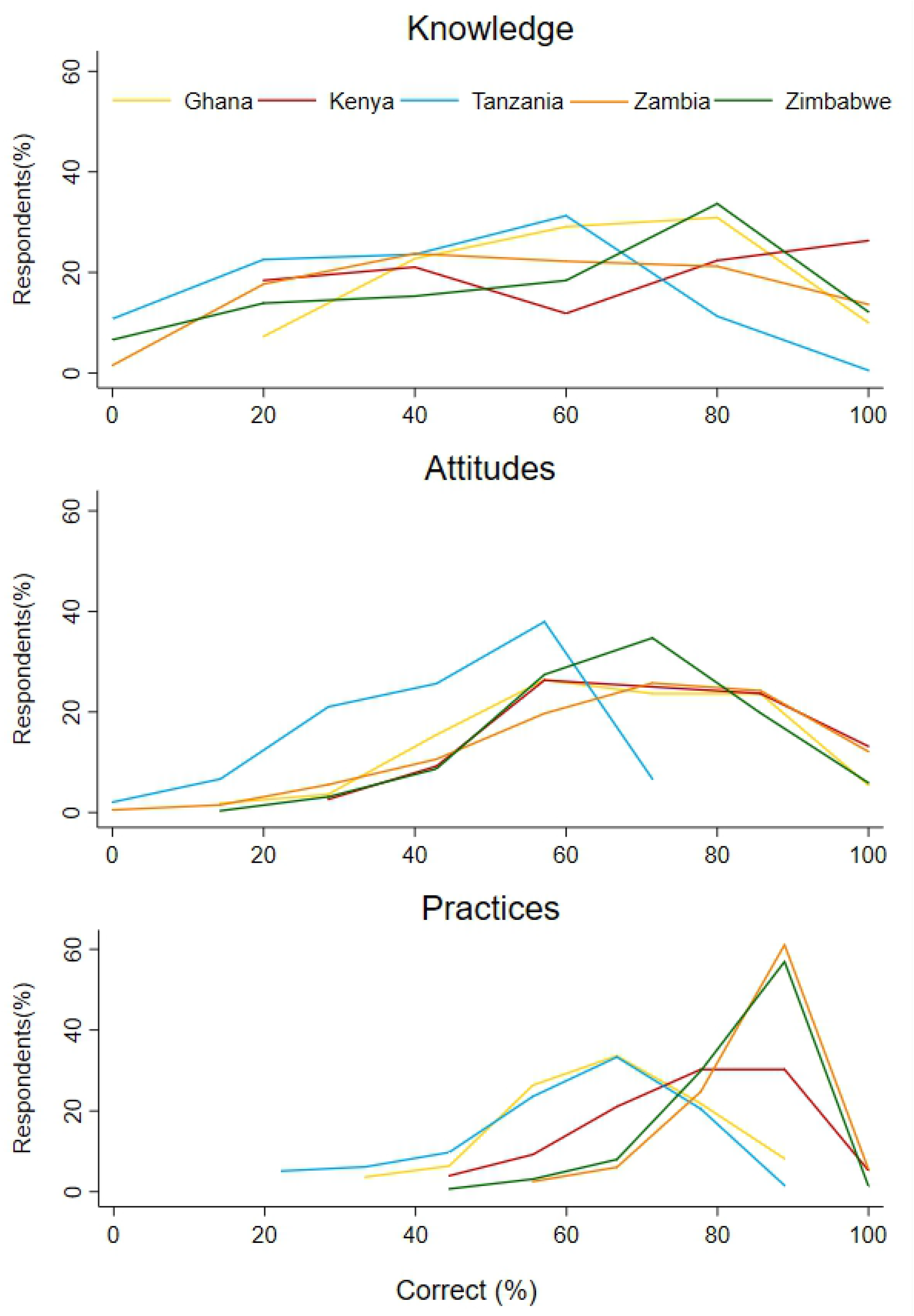
The distribution of scores on AMU/AMR knowledge, attitudes, and practices scales across project countries.

### Associations between KAP across countries

Across countries, KAP scales were all significantly (p<0.05) and positively correlated with the strongest correlations between knowledge and attitudes (≈0.36) and attitudes and practices (≈0.30) (see Table 2). Knowledge and attitudes were also positively and significantly related within countries, ranging from ≈0.21 in Zimbabwe to ≈0.38 in Zambia. In contrast, knowledge was only significantly related to practices in Zambia (≈0.20). Likewise, attitudes were only significantly related to practices in Zambia (≈0.21) and Tanzania, although the latter in the opposite direction (≈-0.15).

**Table 2.**
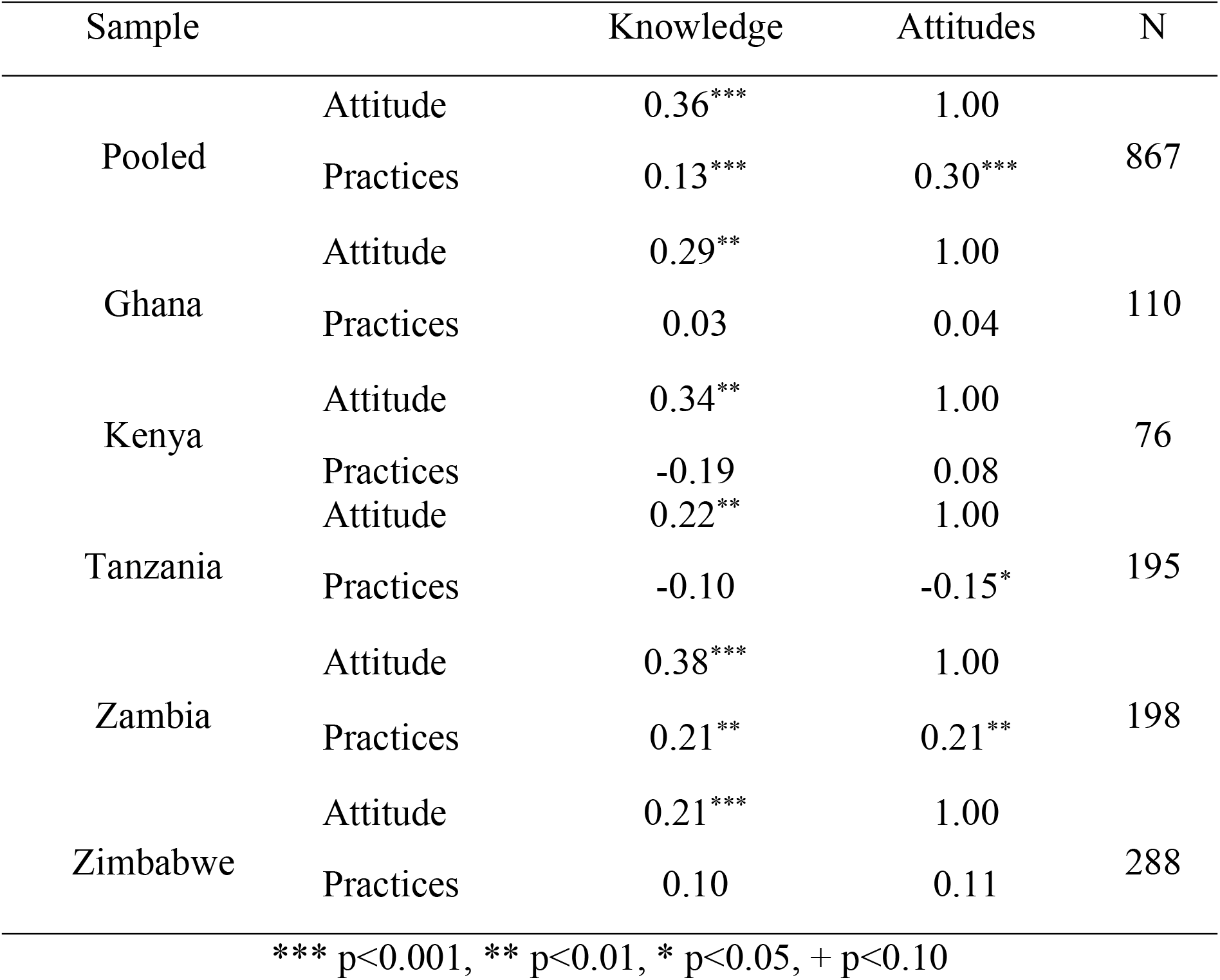
Pearson’s Correlation between KAP measures across and within countries.

### KAP and demographics

Few demographic variables were significantly associated with KAP measures across countries, (Table 3) or poultry and pastoralist systems (Tables S10-S11). Across countries, education level was positively associated with all KAP measures with a one unit increase in education level (i.e., none to primary, primary to secondary, secondary to tertiary) associated with a ≈5% increase in knowledge, a ≈ 3% increase in attitudes, and a ≈ 1% increase in prudent practices. The association between education and KAP indices remained in poultry systems but, in pastoralist systems, education was only significantly associated with knowledge. In all models, after controlling for the impacts of demographics on KAP, there remained significant differences between countries. When country indicators were removed from the models (Table S12), the variance accounted for by the models decreased by approximately a third for knowledge (0.09 vs 0.06) and attitudes (0.24 vs 0.15) and a half for practices (0.42 vs 0.22).

**Table 3.** Associations between KAP measures and demographics. See variable definitions below the table.

| VARIABLES | Knowledge | Attitude | Practices |
| --- | --- | --- | --- |
| *Determinants:* |  |  |  |
| Gender (1=Female) | -0.036*<br>(-0.079 - 0.007) | 0.002<br>(-0.027 - 0.031) | 0.005<br>(-0.014 - 0.023) |
| Age | 0.000<br>(-0.001 - 0.001) | 0.001*<br>(-0.000 - 0.002) | 0.000<br>(-0.000 - 0.001) |
| Education level* | 0.048***<br>(0.023 - 0.073) | 0.027***<br>(0.010 - 0.044) | 0.010*<br>(-0.001 - 0.021) |
| *Country Controls:* |  |  |  |
| Kenya | 0.000<br>(-0.081 - 0.082) | 0.035<br>(-0.020 - 0.091) | 0.102***<br>(0.067 - 0.138) |
| Tanzania | -0.160***<br>(-0.230 - -0.090) | -0.190***<br>(-0.238 - -0.143) | -0.045***<br>(-0.076 - -0.015) |
| Zambia | -0.082**<br>(-0.148 - -0.016) | 0.010<br>(-0.035 - 0.054) | 0.181***<br>(0.152 - 0.209) |
| Zimbabwe | -0.061*<br>(-0.124 - 0.001) | 0.002<br>(-0.040 - 0.044) | 0.162***<br>(0.135 - 0.190) |
| Constant | 0.555***<br>(0.471 - 0.638) | 0.581***<br>(0.525 - 0.638) | 0.632***<br>(0.595 - 0.668) |
| Observations | 866 | 866 | 866 |
| R-squared | 0.092 | 0.246 | 0.428 |

ci in parentheses
\*\*\* p<0.01, \*\* p<0.05, \* p<0.10, + p<0.20
- Education level was recoded into four categories including none=0, primary =1, secondary=2, and tertiary and above=3. Tertiary and above indicates any additional education after secondary school, including certificates, diplomas, bachelors, masters, and PhDs

### KAP and on-farm dynamics

Whether a respondent observed treatment failure was positively associated with high level of knowledge (4% increase) while it was negatively associated with good practices (3% decrease) (Table 4). Likewise, while disease level was positively associated with attitudes, with every additional disease predicting a ≈15% increase in prudent attitude scores, disease levels were negatively associated with prudent practices with every unit increase associated with a ≈17% decrease in prudent practices. Having training in farm management was significantly associated with better knowledge (≈17% increase) and more prudent attitudes (≈3% increase) but a decrease in prudent practices (≈3%). Number of AMs was positively associated with knowledge with a 0.2 % increase for every additional AM reportedly used. The significant associations between KAP scales and training and the number of AMs reportedly used were maintained in the poultry model. In addition, whether a household kept records was positively associated with knowledge and attitude scores, with respondents reporting to keep records expected to have ≈10% greater knowledge and ≈3% more prudent attitudes. Disease level was not related to KAP scales in poultry systems (Table S13). In pastoralist systems, disease levels were the only significant farm characteristic related to KAP, with every additional disease predicting a ≈35% increase in prudent attitudes and a ≈44% decrease in prudent practices (Table S14).

**Table 4.**
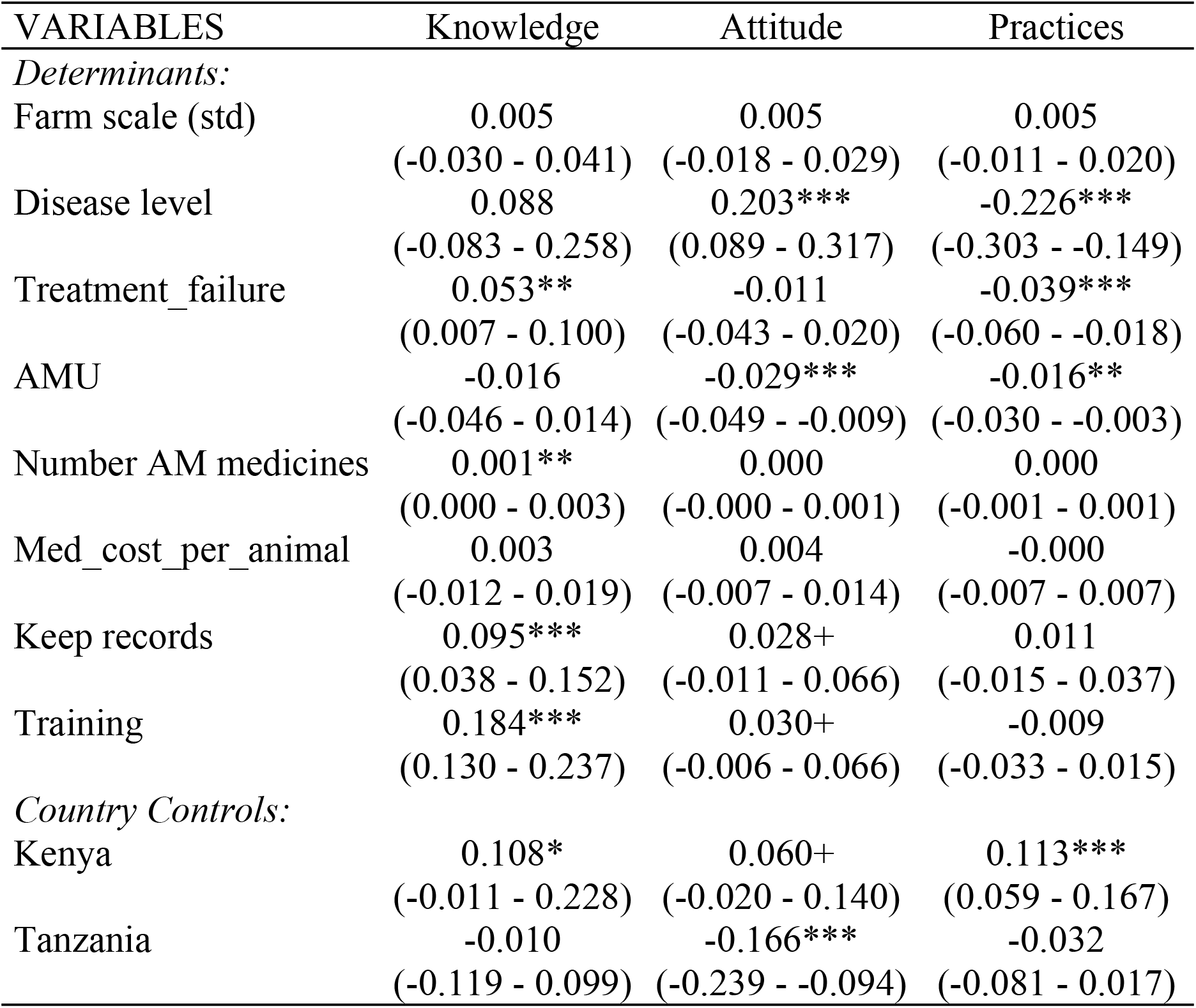
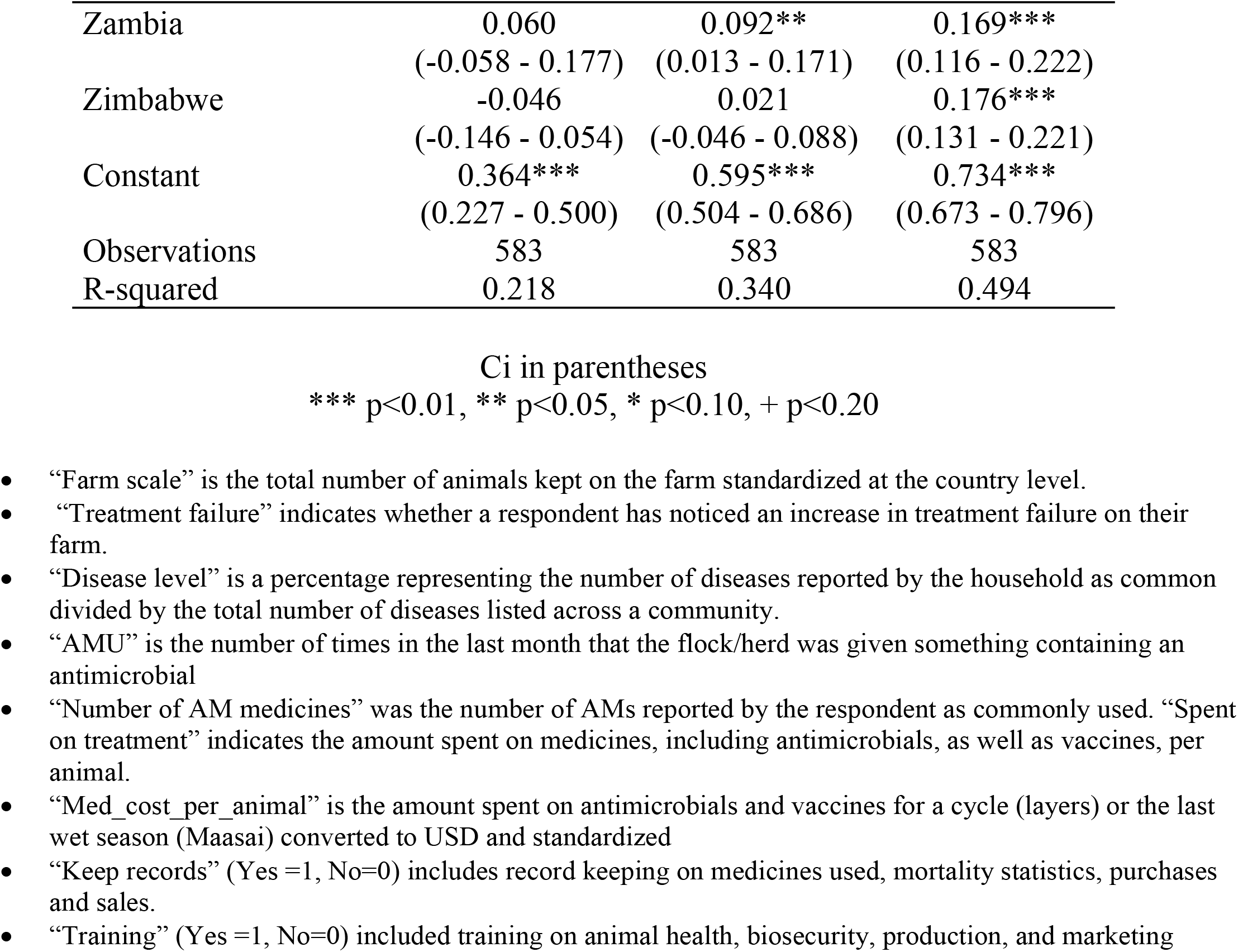
Associations between farm characteristics and KAP constructs adjusted for country effects. See variable definitions below the table.

### KAP and sources of animal health advice

Across countries, AMU/AMR knowledge was positively associated with seeking health advice from extension workers and government veterinarians (Table 5). Each increase in frequency of seeking advice from extension workers and government veterinarians, (i.e., never to sometimes, sometimes to almost always), increased knowledge scores by ≈6% and ≈7%, respectively. Frequency of advice from extension and government workers was also associated with prudent attitudes with every increase in advice frequency associated with around a ≈3% increase. In contrast, while the frequency of seeking advice from government veterinarians was associated with a ≈2% increase in prudent practices, seeking advice from extension workers was negatively associated with practices (≈3% decrease). Frequency of getting advice from agrovets did not significantly impact any KAP measures, but the type of information provided in agrovet interactions (i.e., instructions on use, what AM to use) was positively associated with more prudent practices (≈2% increase). No other source of animal health information was significantly related to KAP measures, including private veterinarians, family/friends, and laboratory workers. In poultry systems, associations between getting advice from extension workers and government veterinarians had similar impacts on KAP scales, although larger in magnitude (Table S15). In addition, seeking health advice from private veterinarians was positively associated with knowledge (≈4% increase) and practices (≈1.4% increase). In pastoralist systems, the sole source of health advice significantly associated with KAP measures was frequency of seeking health advice from extension workers, which was associated with a 5% increase in attitudes (Table S16). Acquiring more information from agrovets was associated with a ≈3% decrease in prudent attitudes but a 9% increase in reported prudent use practices.

**Table 6.**
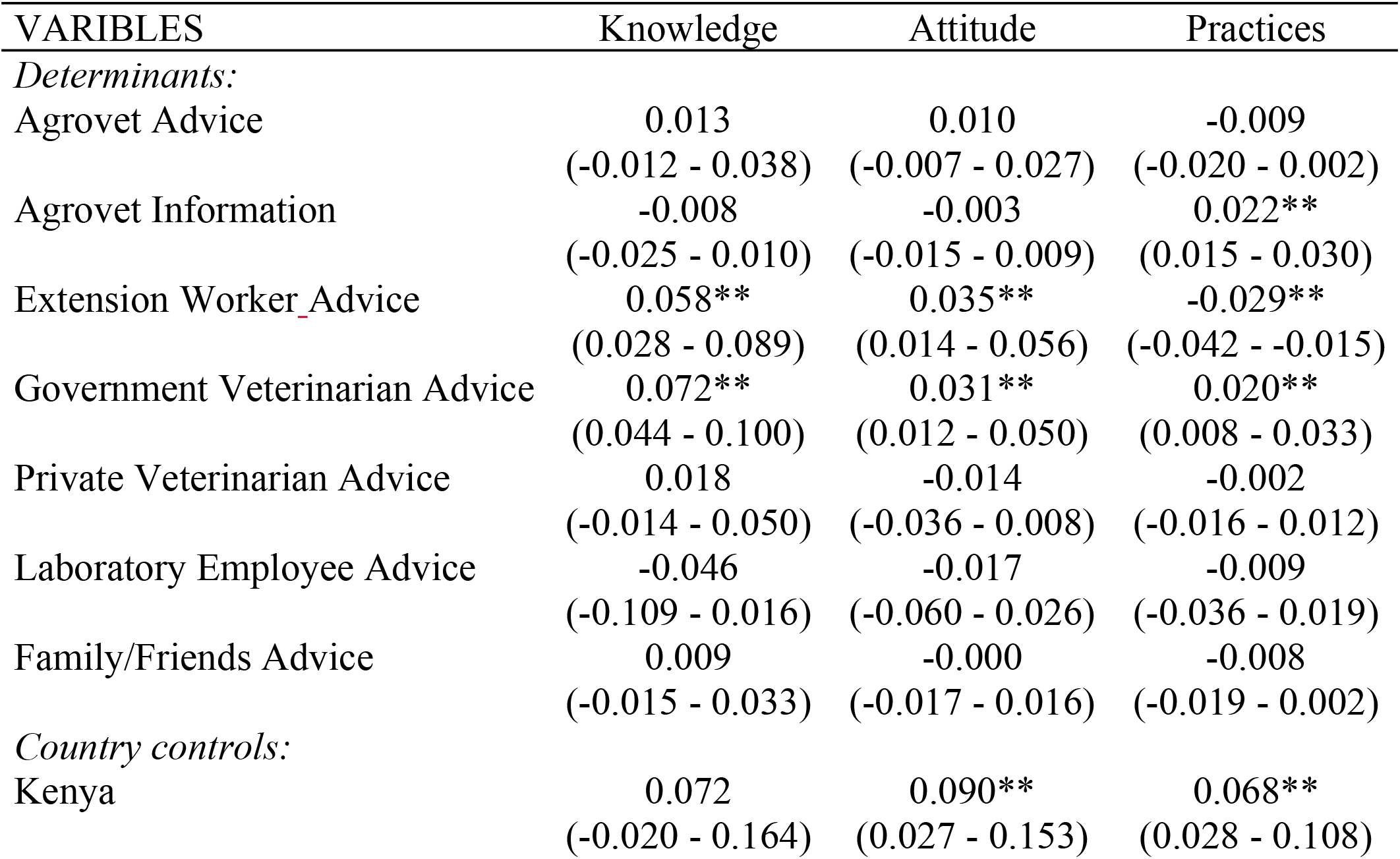
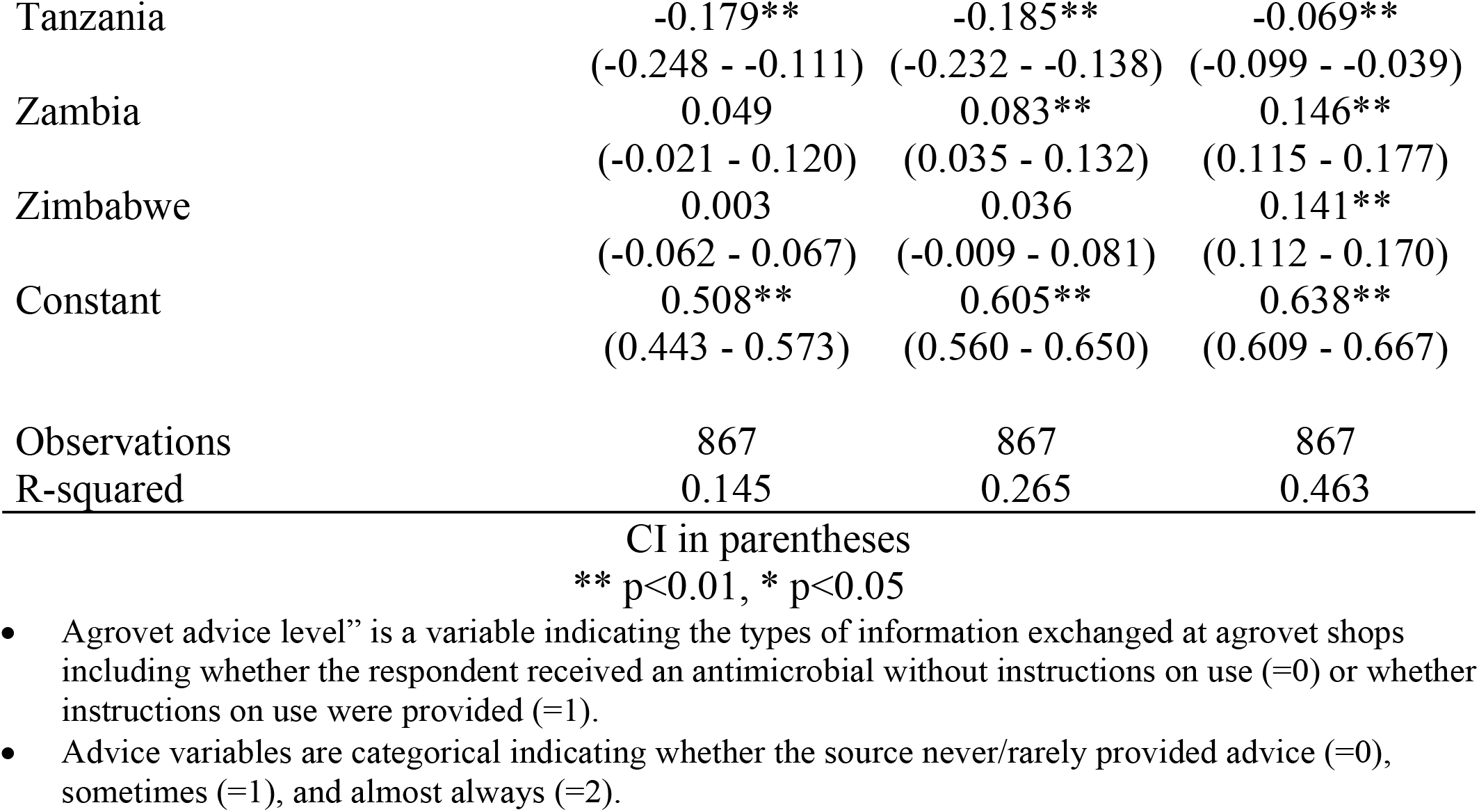
Correlates between KAP measures and sources of animal health advice. See variable definitions below the table.

## Discussion

In this study we examined the knowledge, attitudes, and practices related to antimicrobial use and resistance in livestock farmers across five African countries (Ghana, Kenya, Tanzania, Zambia, Zimbabwe). Broadly, we found that it is individuals who live or work at the farm who draw upon their knowledge and experiences to make decisions regarding animal healthcare, most often with input from family, friends and neighbors. It is also these individuals who are responsible for administering antimicrobials to their animals, usually for therapeutic reasons in poultry systems and more often for growth and prevention in Maasai pastoralists. Reliance on animal health professionals for advice and treatment is limited with only around 10% of respondents regularly relying upon veterinarians for advice and 5% for treatment. A consequence of this limited interaction is clear in prescription practices, with just 12% of the 867 farmers surveyed reporting having a prescription when purchasing antimicrobial products. When purchasing these products, over 83% of respondents said they relied upon local agrovet drug shops. Few farmers reported observing withdrawal periods after treatment with antimicrobials, both consuming and selling animal products (meat, milk, eggs) from animals currently undergoing treatment or still within the withdrawal period.

High levels of engagement in the “informal veterinary sector” – characterized by “lay” diagnoses and treatment, limited input from trained health professionals, and the practicing of other non-prudent practices (e.g., use of nonprescribed antimicrobials, nonadherence to withdrawal periods)– are consistent with findings from other studies conducted in the project countries including among Maasai pastoralists in Tanzania and Kenya [26,30,42,43], cattle keepers in Eastern Province, Zambia [44] and poultry farmers in Ghana, Kenya, Tanzania, Zambia, and Zimbabwe [26–35] (see Table S17 for a more details of these studies). Collectively, these results further confirm the negative impacts of the Structural Adjustment Policies that relegated the public veterinary services in many African countries to policy and regulation functions [45]. The few veterinarians who went into private practice were often located in areas where they could make good returns, which often meant they were removed from farmers in rural areas, especially those inhabiting arid environments such as pastoralists. As our results suggest, this shift towards paid veterinary services has continued to limit access to animal health professionals with voids in veterinary care being occupied by persons operating as agrovets. These individuals, while doubling as drug shop agents and service providers, often having limited levels of formal training in animal health and knowledge on AMU/AMR.

Limited access to animal health professionals and engagement in the informal veterinary sector holds several consequences for National Action Plans to address AMR in LMICs. Namely, intervention strategies to motivate prudent AMU and limit AMR must recognize the necessity of contextualizing antimicrobial use and related practices at the farm level. This contextualization will represent a bottom-up understanding of the sociocultural, economic, and historical and professional factors driving patterns of AMU/AMR knowledge, attitudes, and practices across different livestock systems and varying production scales. Our results highlight that these factors are likely to vary across cultures. For example, farmers in poultry systems could recount significantly greater amounts of AMU/AMR knowledge and reported more prudent attitudes and practices compared to Maasai pastoralists. These differences may be due to varying educational backgrounds as poultry farmers tended to be better educated and we found a positive association between education level and KAP scales. These differences could also be driven by differences in indigenous ethnoveterinary belief systems, which the Maasai draw heavily upon to diagnosis and treat their animals. If these beliefs are inconsistent with concepts underlying AMR (e.g., selection, transmission), then some Maasai may not perceive AMR as risky or believe AMU could adversely impact their livelihoods [46]. Establishing the relative contributions of the different livelihood factors on knowledge, attitudes and practices related to AMU and AMR will be critical in resource-limited contexts such as LMICs where only a limited number of factors can be targeted within behavioral change interventions.

Our results stress that intervention philosophies based primarily upon top-down approaches are unlikely to result in broad changes in AMU within LMICs. Instead of pursuing blanket top-down strategies, a more effective approach to address AMU and AMR in LMICs would be to develop smaller scale behavioral change interventions targeted at stakeholders along the chain of antimicrobial use. Our results, for example, show that agrovet employees are often the only stakeholders outside the household with an animal health background that farmers interact with between deciding on treatment and administering antimicrobials. Our regression models showed that respondents who were given more information from these agrovets (i.e., advice on recommended antimicrobials and treatment regimes) also reported more prudent practices (i.e., 2% higher across all systems, 3% in poultry and 8% in pastoralist). These findings suggest that interventions aimed at leveraging the influence of agrovets may be able to impact local AMU patterns. However, given the variability in training among agrovet employees [43,52], a better understanding of the knowledge, attitudes, and practices of agrovets will be needed to develop these interventions.

### Study Limitations and Future Directions

This study represents one of the most comprehensive surveys of AMU/AMR-related KAP across livestock systems in Africa, but more robust ethnographic work is needed across cultures and production systems and scales to identify the factors associated with KAP variation. While we found that several demographic, livelihood and health-seeking factors were significantly related to KAP measures, these factors usually accounted for low levels of variance (<5%). Most of the variance was accounted for by country indicators, especially for practices, which significantly varied across countries and clearly hold the greatest consequences for the distribution of AMU and AMR. The contribution of country-level indicators suggests the existence of factors (cultural, historical, economic) that were not, or could not be, recovered in our mostly self-report cross-sectional surveys. Identification of some of these factors will require different study designs (i.e., longitudinal) and ethnographic techniques (participant-observation). Observational methods, for example, are warranted given evidence of self-desirability bias in the reporting of prudent practices. Indeed, while our focus group participants confessed to not following withdrawal periods, our KAP results indicated that up to 30% of households reported observing withdrawal periods in some countries. Recall biases also impact the collection of antimicrobial use data. Methods such as repeated self-report measures, passive surveillance methods, including collection of used sachets and bottles in in waste buckets, and triangulation with sales data at agrovets will be needed to produce accurate and quantitative antimicrobial use data. Intervention strategies must also reconcile the lack of association between practices and knowledge and attitudes in most countries. Indeed, especially in poultry systems, the observance of non-prudent practices cannot be explained by a lack of knowledge or attitudes concerning AMR. Around 70% of respondents held sufficient knowledge to understand the AMR issue and associated desirable attitudes.

Finally, while our results emphasize the importance of bottom-up strategies to address AMU and AMR there remains a clear need for broader infrastructural changes in LMICs that can support implementation of top-down strategies (e.g., increase in the veterinary workforce, better resourced food safety departments) [53]. Our models of health-seeking practices, for example, demonstrated the positive role that the professional veterinary sector can play in promoting prudent AMU practices. Farmers who more frequently sought advice from professional veterinarians reported more prudent knowledge and attitudes (6%-7% increases), and better practices (2%-3% increases). Considering these trends, intervention strategies targeted at veterinarians (e.g., stewardship programs) *could* prove successful if they were coupled with broader infrastructural changes to increase access to animal health professionals. Importantly, while top-down strategies are needed, they must still be informed by the realities of AMU/AMR or they may risk doing more harm than good. Wholesale application of some popular top-down regulations in LMICs (restrictions on certain types of antimicorbials) risks preventing access to antimicrobials in contexts where a lack of access to these drugs continues to result in the deaths of considerably more people and animals than resistant infections [4,24].

## Conclusion

This study has shown that livestock farmers in five African countries make decisions on antimicrobial with limited inputs from animal health professionals, who are often inaccessible due to distance and/or financial considerations. Given these realities, interventions to promote prudent use practices in the short-term must be focused on the contexts where animal healthcare decisions are made–at the farm level– and be delivered using channels that have been demonstrated to be impactful. Until the livestock sectors within these countries can attract qualified veterinarians to provide quality services including better AMU, fellow community members and agrovet shop workers will continue to provide veterinary advice and care. Therefore, any interventions aimed at optimizing AMU must include these key stakeholders. These interventions should be founded upon a bottom-up approach that documents the KAP driving AMU, while also identifying the factors associated with KAP. Conducting these studies across cultural and production contexts will provide the much-needed evidence base to develop and implement targeted behavioral change interventions to reduce AMR globally.

## References

1. Davies J, Davies D. Origins and Evolution of Antibiotic Resistance. Microbiol Mol Biol Rev. 2010;74: 417–433. doi:10.1128/MMBR.00016-10

2. Landers TF, Cohen B, Wittum TE, Larson EL. A Review of Antibiotic Use in Food Animals: Perspective, Policy, and Potential. Public Health Reports. 2012;127: 4–22.

3. Levy SB, Marshall B. Antibacterial resistance worldwide: causes, challenges and responses. Nature Medicine. 2004;10: S122–S129. doi:10.1038/nm1145

4. Palmer GH, Call DR. Antimicrobial resistance: A global public health challenge requiring a global one health strategy [Internet]. 2013. Available: https://www.iom.edu/~/media/Files/Perspectives-Files/2013/Commentaries/BGH-Anitmicrobial-Resistance.pdf

5. World Health Organization. Global action plan on antimicrobial resistance. Geneva, Switzerland: World Health Organization; 2015.

6. Van Boeckel TP, Brower C, Gilbert M, Grenfell BT, Levin SA, Robinson TP, et al. Global trends in antimicrobial use in food animals. Proc Natl Acad Sci USA. 2015;112: 5649. doi:10.1073/pnas.1503141112

7. Robinson TP, Bu DP, Carrique-Mas J, Fèvre EM, Gilbert M, Grace D, et al. Antibiotic resistance is the quintessential One Health issue. Transactions of the Royal Society of Tropical Medicine and Hygiene. 2016;110: 377–380. doi:10.1093/trstmh/trw048

8. Guenther S, Ewers C, Wieler L. Extended-Spectrum Beta-Lactamases Producing E. coli in Wildlife, yet Another Form of Environmental Pollution? Frontiers in Microbiology. 2011;2: 246. doi:10.3389/fmicb.2011.00246

9. Katakweba A, Møller K, Muumba J, Muhairwa A, Damborg P, Rosenkrantz JT, et al. Antimicrobial resistance in faecal samples from buffalo, wildebeest and zebra grazing together with and without cattle in T anzania. Journal of Applied microbiology. 2015;118: 966–975.

10. Afema JA, Byarugaba DK, Shah DH, Atukwase E, Nambi M, Sischo WM. Potential Sources and Transmission of Salmonella and Antimicrobial Resistance in Kampala, Uganda. PLOS ONE. 2016;11: e0152130. doi:10.1371/journal.pone.0152130

11. Mather A, Reid S, Maskell D, Parkhill J, Fookes M, Harris S, et al. Distinguishable epidemics of multidrug-resistant Salmonella Typhimurium DT104 in different hosts. Science. 2013;341: 1514–1517.

12. Mather AE, Matthews L, Mellor DJ, Reeve R, Denwood MJ, Boerlin P, et al. An ecological approach to assessing the epidemiology of antimicrobial resistance in animal and human populations. Proc Biol Sci. 2011; doi:10.1098/rspb.2011.1975

13. Dorado-García A, Smid JH, van Pelt W, Bonten MJM, Fluit AC, van den Bunt G, et al. Molecular relatedness of ESBL/AmpC-producing Escherichia coli from humans, animals, food and the environment: a pooled analysis. Journal of Antimicrobial Chemotherapy. 2018;73: 339–347. doi:10.1093/jac/dkx397

14. Hendriksen RS, Munk P, Njage P, van Bunnik B, McNally L, Lukjancenko O, et al. Global monitoring of antimicrobial resistance based on metagenomics analyses of urban sewage. Nature Communications. 2019;10: 1124. doi:10.1038/s41467-019-08853-3

15. Collignon P, Beggs JJ, Walsh TR, Gandra S, Laxminarayan R. Anthropological and socioeconomic factors contributing to global antimicrobial resistance: a univariate and multivariable analysis. The Lancet Planetary Health. 2018;2: e398–e405. doi:10.1016/S2542-5196(18)30186-4

16. Dorado-García A, Dohmen W, Bos MEH, Verstappen KM, Houben M, Wagenaar JA, et al. Dose-Response Relationship between Antimicrobial Drugs and Livestock-Associated MRSA in Pig Farming1. Emerging Infectious Diseases. 2015;21: 950–959. doi:10.3201/eid2106.140706

17. Bebell LM, Muiru AN. Antibiotic use and emerging resistance: how can resource-limited countries turn the tide? Global heart. 2014;9: 347–358. doi:10.1016/j.gheart.2014.08.009

18. Cox JA, Vlieghe E, Mendelson M, Wertheim H, Ndegwa L, Villegas MV, et al. Antibiotic stewardship in low- and middle-income countries: the same but different? Clinical Microbiology and Infection. 2017;23: 812–818. doi:10.1016/j.cmi.2017.07.010

19. Founou LL, Founou RC, Essack SY. Antibiotic Resistance in the Food Chain: A Developing Country-Perspective. Frontiers in microbiology. 2016;7: 1881–1881. doi:10.3389/fmicb.2016.01881

20. Okeke IN, Laxminarayan R, Bhutta ZA, Duse AG, Jenkins P, O’Brien TF, et al. Antimicrobial resistance in developing countries. Part I: recent trends and current status. The Lancet Infectious Diseases. 2005;5: 481–493. doi:10.1016/S1473-3099(05)70189-4

21. Food and Agriculture Organization. Monitoring global progress on addressing antimicrobial resistance: analysis report of the second round of results of AMR country self-assessment survey 2018 [Internet]. Rome, Italy: Food and Agriculture Organization; 2018. Available: http://www.fao.org/publications/card/en/c/CA0486EN

22. World Health Organization. Antimicrobial resistance: a manual for developing national action plans. 2016;

23. Gauthier J, Simeon M, De Haan C. The effect of structural adjustment programmes on the delivery of veterinary services in Africa. Citeseer; 1999. pp. 133–156.

24. Sones KR, Catley A, editors. Primary animal health care in the 21st Century: shaping the rules, policies and institutions. Proceedings of an international conference held in Mombaa, Kenya, 15-18 October 2002. Nairobi, Kenya: African/Union/Interafrican Bureau for Animal Resources; 2003.

25. Omulo S, Thumbi SM, Njenga MK, Call DR. A review of 40 years of enteric antimicrobial resistance research in Eastern Africa: what can be done better? Antimicrobial Resistance and Infection Control. 2015;4: 1–13. doi:10.1186/s13756-014-0041-4

26. Caudell MA, Quinlan MB, Subbiah M, Call DR, Roulette CJ, Roulette JW, et al. Antimicrobial Use and Veterinary Care among Agro-Pastoralists in Northern Tanzania. PloS one. 2017;12: e0170328.

27. Johnson S, Bugyei K, Nortey P, Tasiame W. Antimicrobial drug usage and poultry production: case study in Ghana. Anim Prod Sci. 2019;59: 177–182.

28. Nonga H, Simon C, Karimuribo E, Mdegela R. Assessment of antimicrobial usage and residues in commercial chicken eggs from smallholder poultry keepers in Morogoro municipality, Tanzania. Zoonoses and Public health. 2010;57: 339–344.

29. Turkson P. Use of drugs and antibiotics in poultry production in Ghana. Ghana Journal of Agricultural Science. 2008;41.

30. Roderick S, Stevenson P, Mwendia C, Okech G. The Use of Trypanocides and Antibiotics by Maasai Pastoralists. Tropical Animal Health and Production. 2000;32: 361–374. doi:10.1023/A:1005277518352

31. Boamah VE, Agyare C, Odoi H, Dalsgaard A. Practices and factors influencing the use of antibiotics in selected poultry farms in Ghana. 2016;

32. Mubito EP, Shahada F, Kimanya ME, Buza JJ. Antimicrobial use in the poultry industry in Dar-es-Salaam, Tanzania and public health implications. Am J Res Commun. 2014;2: 51–63.

33. Gehring R, Swan G, Sykes R. Supply of veterinary medicinal products to an emerging farming community in the North West Province of South Africa. Journal of the South African Veterinary Association. 2002;73: 185–189.

34. Ojo OE, Fabusoro E, Majasan AA, Dipeolu MA. Antimicrobials in animal production: usage and practices among livestock farmers in Oyo and Kaduna States of Nigeria. Tropical animal health and production. 2016;48: 189–197.

35. Auta A, Hadi MA, Oga E, Adewuyi EO, Abdu-Aguye SN, Adeloye D, et al. Global access to antibiotics without prescription in community pharmacies: A systematic review and meta-analysis. Journal of Infection. 2018;

36. Creswell JW, Clark VLP. Designing and conducting mixed methods research. Third. Thousand Oaks, CA: Sage publications; 2017.

37. Guest G, Namey E, McKenna K. How many focus groups are enough? Building an evidence base for nonprobability sample sizes. Field methods. 2017;29: 3–22.

38. Neter J, Kutner MH, Nachtsheim CJ, Wasserman W. Applied linear statistical models. Chicago: Irwin Chicago; 1996.

39. O’Brien RM. Dropping highly collinear variables from a model: why it typically is not a good idea. Social Science Quarterly. 2017;98: 360–375.

40. Cohen P, Cohen J, West S, Aiken L. Applied Multiple Regression/Correlation Analysis for the Behavioral Sciences. Mahwah, New Jersey: Lawrence Erlbaum Associates; 2003.

41. Huber PJ. The behavior of maximum likelihood estimates under nonstandard conditions. University of California Press; 1967. pp. 221–233.

42. Ngumbi AF, Silayo RS. A cross-sectional study on the use and misuse of trypanocides in selected pastoral and agropastoral areas of eastern and northeastern Tanzania. Parasites & vectors. 2017;10: 607.

43. Higham LE, Ongeri W, Asena K, Thrusfield MV. Characterising and comparing drug-dispensing practices at animal health outlets in the Rift Valley, Kenya: an exploratory analysis (part II). Tropical Animal Health and Production. 2016;48: 1633–1643. doi:10.1007/s11250-016-1137-z

44. Delespaux V, Geerts S, Brandt J, Elyn R, Eisler M. Monitoring the correct use of isometamidium by farmers and veterinary assistants in Eastern Province of Zambia using the isometamidium-ELISA. Veterinary parasitology. 2002;110: 117–122.

45. The veterinary pharmaceutical industry in Africa: a study of Kenya, Uganda and South Africa. Nairobi, Kenya: African Union/ Interafrican Bureau for Animal Resources (AU/IBAR); 2004.

46. Caudell MA, Quinlan MB, Quinlan RJ, Call DR. Medical pluralism and livestock health: ethnomedical and biomedical veterinary knowledge among East African agropastoralists. Journal of Ethnobiology and Ethnomedicine. 2017;13: 7–15.

47. Speksnijder DC, Mevius DJ, Bruschke CJM, Wagenaar JA. Reduction of Veterinary Antimicrobial Use in the Netherlands. The Dutch Success Model. Zoonoses and Public Health. 2014;62: 79–87. doi:10.1111/zph.12167

48. Food and Agriculture Organization of the United Nations (FAO). Tackling antimicrobial use and resistance in pig production: lessons learned in Denmark [Internet]. Rome, Italy: FAO; 2019 Jun p. 55. Available: http://www.fao.org/3/ca2899en/CA2899EN.pdf

49. Dupont N, Diness LH, Fertner M, Kristensen CS, Stege H. Antimicrobial reduction measures applied in Danish pig herds following the introduction of the “Yellow Card” antimicrobial scheme. Preventive Veterinary Medicine. 2017;138: 9–16. doi:10.1016/j.prevetmed.2016.12.019

50. Kalungia AC, Burger J, Godman B, Costa J de O, Simuwelu C. Non-prescription sale and dispensing of antibiotics in community pharmacies in Zambia. Expert review of anti-infective therapy. 2016;14: 1215–1223.

51. Mukokinya MMA, Opanga S, Oluka M, Godman B. Dispensing of antimicrobials in Kenya: A cross-sectional pilot study and its implications. Journal of research in pharmacy practice. 2018;7: 77.

52. Bett B, Machila N, Gathura PB, McDemott JJ, Eisler MC. Characterisation of shops selling veterinary medicines in a tsetse-infested area of Kenya. Preventive Veterinary Medicine. 2004;63: 29–38. doi: 10.1016/j.prevetmed.2004.02.004

53. Caudell MA, Mair C, Subbiah M, Matthews L, Quinlan RJ, Quinlan MB, et al. Identification of risk factors associated with carriage of resistant Escherichia coli in three culturally diverse ethnic groups in Tanzania: a biological and socioeconomic analysis. The Lancet Planetary Health. 2018;2: e489–e497. doi:10.1016/S2542-5196(18)30225-0

